# The alopecia areata phenotype is induced by the water avoidance stress test in *cchcr1*-deficient mice

**DOI:** 10.1101/2020.12.16.423031

**Authors:** Qiao-Feng Zhao, Nagisa Yoshihara, Atsushi Takagi, Etsuko Komiyama, Akira Oka, Shigaku Ikeda

**Author notes:** Corresponding author: Shigaku Ikeda, MD, PhD, Department of Dermatology and Allergology, Juntendo University Graduate School of Medicine, 2-1-1 Hongo, Bunkyo-ku, Tokyo 113-8421, Japan. COI: The authors have no conflicts of interest to declare.

## Abstract

**Background:** We recently discovered a nonsynonymous variant in the coiled-coil alpha-helical rod protein 1 (*CCHCR1*) gene within the alopecia areata (AA) risk haplotype; mice engineered to carry the risk allele displayed a hair loss phenotype.

**Objective:** To further investigate the involvement of the *CCHCR1* gene in AA pathogenesis.

**Methods:** We developed an AA model using C57BL/6N *cchcr1* gene knockout mice. Mice (6-8 weeks) were divided into two groups: *cchcr1* ^-/-^ mice and wild-type (WT) mice. Both groups were subjected to a water avoidance stress (WAS) test.

**Results:** Eight weeks after the WAS test, 25% of *cchcr1* mice exhibited noninflammatory foci of alopecia on the dorsal skin. The foci resembled human AA in terms of gross morphology, trichoscopic findings and histological findings.

**Conclusions:** Our results strongly suggest that *CCHCR1* is associated with AA pathogenesis and that *cchcr1*^-/-^ mice are a good model for investigating AA.

**Author summary:** Alopecia areata is thought to affect 1-2% of the population. In severe alopecia areata, changes in appearance significantly reduce the patient’s quality of life, but there is no established treatment. Its pathogenic mechanism is thought to cause an autoimmune disease of the hair bulb of growing hair. Many genes and factors associated with AA onset form a background that easily causes such an immune reaction, and the causative genes related to alopecia areata are being elucidated worldwide. We were able to identify the *CCHCR1* gene as one of the causative genes by genome analysis. In this study, we created *CCHCR1*-deficient mice using Cre/loxP technology and confirmed that 25% of *CCHCR1*-deficient mice that underwent the WAS test for psychological stimulation developed hair loss similar to that observed in human alopecia areata. This suggests that the *CCHCR1* gene is a disease susceptibility gene for alopecia areata.

## Introduction

Alopecia areata (AA) is a complex genetic and tissue-specific autoimmune disease characterized by nonscarring hair loss that may begin as patches that can coalesce and progress to cover the entire scalp and/or whole body [1]. Most authors tend to accept the hypothesis that AA is caused by a T-cell mediated autoimmune response targeting an unknown antigen in anagen-stage hair follicles [2,3].

Previous genome-wide association studies (GWAS) have implicated a number of immune and nonimmune loci in the etiology of AA, although none have yet been demonstrated to be causative for the disease, and none have been functionally confirmed to be involved in AA pathogenesis [4,5]. Alleles of the human leukocyte antigen (HLA) genes within the major histocompatibility complex (MHC) on chromosome 6p21.3 have so far shown the strongest associations with AA across different ethnic groups [4,5]. The largest reported genome-wide meta-analysis of AA identified HLA-DRβ1 as a key etiologic driver [4]. However, no convincing susceptibility gene has yet been pinpointed in the MHC.

We recently discovered a nonsynonymous variant (rs142986308, p. Arg587Trp) in the MHC associated with AA susceptibility in *CCHCR1* (coiled-coil alpha-helical rod protein 1), which encodes a novel component of hair shafts. In addition, our results demonstrate that mice carrying this amino acid substitution display a patchy hair loss phenotype. We further identified keratin abnormalities in the hair shaft and comparative differential expression of hair-related keratin genes not only in alopecic mice but also in hair follicles from AA patients with the risk variant. Thus, our study identified a novel AA susceptibility variant that was validated by functional analysis [6].

The purpose of this study was to investigate the effect of *CCHCR1* gene deficiency on AA pathogenesis. First, we generated C57BL/6N mice with deletion of the *cchcr1* gene using Cre/loxP technology. Because *cchcr1*^−/−^ mice develop normally and exhibit no obvious behavioral or physical phenotypic defects and because psychological stress is known to be a triggering factor in AA patients [7], the water avoidance stress (WAS) test, which is well known to stimulate psychological stress in mice [8,9], was performed on *cchcr1*^*−/−*^ and WT mice.

## Results

First, we generated mice with deletion of *cchcr1* using Cre/loxP technology. The final recombined floxed allele is presented in Fig. 1a. *Cchcr1*^−/−^ mice were obtained by intercrossing heterozygous (*cchcr1* ^*+/−*^) mice. After identifying Cre-mediated deletion and the inactivation of *cchcr1*, the expression of *cchcr1* was assessed by PCR (Fig. 1b). We also analyzed the protein and RNA levels of *cchcr1* in the skin (Fig. 1c, 1d). The *cchcr1* ^*−/−*^ mice were born at the expected ratios and had normal weights. The mice developed normally, were fertile, and exhibited no obvious behavioral or physical phenotypic defects.

**Fig 1.**
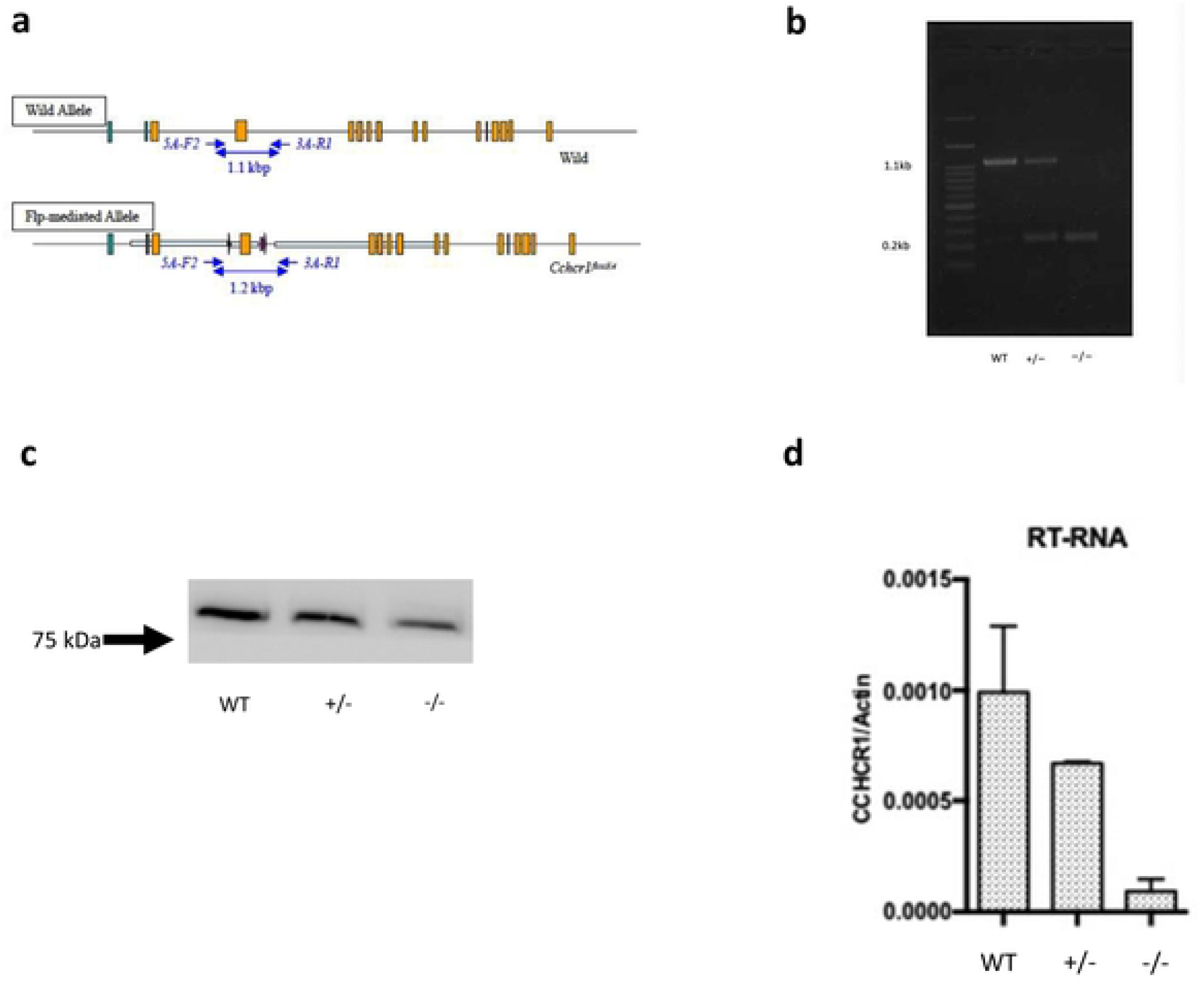
**Fig 1a.** Exon 4 was deleted by using the loxP site in *cchcr1*. The mice were backcrossed with wild-type (WT) C57BL/6N mice. Homozygous (*cchcr1*^*−/−*^) knockout (KO) mice were obtained by intercrossing heterozygous (*cchcr1* ^+*/−*^) mice, and then genotyping was performed by PCR. **Fig 1b.** Genotypes were identified by PCR using tail tips. Genotyping confirmed the presence of WT mice, *cchcr1*^+*/−*^ mice and *cchcr1*^*−/−*^ mice. The samples were loaded as follows: first lane, molecular weight marker; the WT band at 1.1 kbp; the two *cchcr1*^+*/−*^ bands at 1.1 kbp and 0.2 kbp; and the *cchcr1* ^*−/−*^ band at 0.2 kbp. **Fig 1c.** Western blot showing the expression of *cchcr1* in skin tissue lysates from WT, *cchcr1*^*+/−*^, and *cchcr1*^*-/-*^ mice. **Fig 1d.** The expression of *cchcr1* in WT mice, *cchcr1*^*+/−*^ mice, and *cchcr1*^*-/-*^ mice was examined by RT-PCR. β-actin was used as an internal control.

Next, a WAS test was performed on *cchcr1*^-/-^ mice and WT mice (n=9) (both 6-8 weeks after birth). The WAS test was performed for two hours a day, five times per week, for two weeks.

Eight weeks after the WAS test, in contrast to WT mice (Fig. 2c), approximately 25% of *cchcr1* ^*-/-*^ mice spontaneously exhibited noninflammatory foci of alopecia in the dorsal area (Fig. 2a, Table 1). Hair loss may also develop within a sharply demarcated, localized area or diffusely on the dorsal side. The skin of the hair loss area has a normal appearance. The hair-pull test showed an increase in dystrophic anagen hairs. Characteristic phenotypes of AA (short stubs of hair and exclamation mark hairs) were observed in the hair loss area by dermoscopy. We also identified white dots and tapering hairs as signs of AA (Fig. 2a).

**Table 1.**
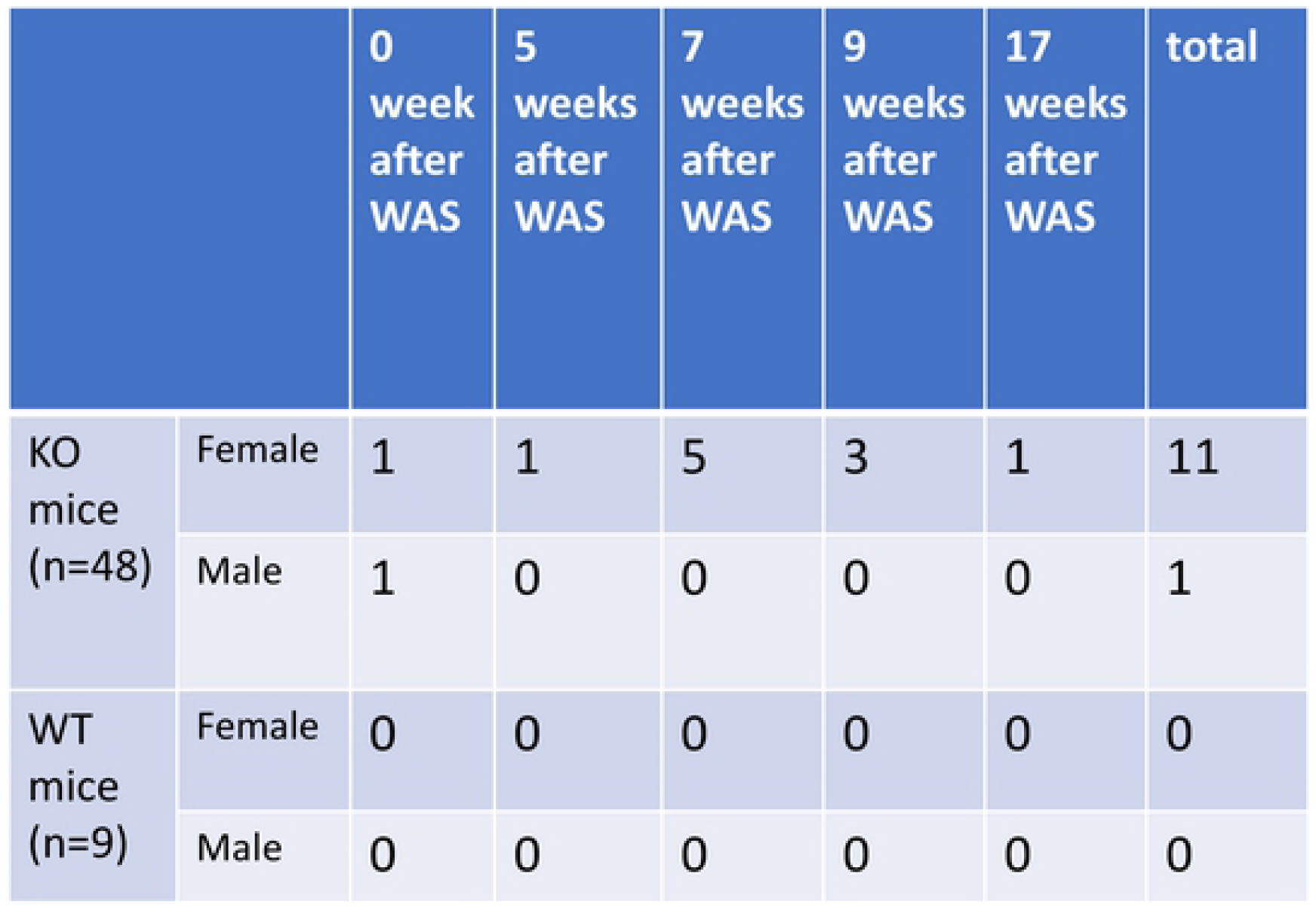
AA-like skin lesions on *cchcr1* ^*-/-*^ mice (n=48) and WT control mice (n=9) after the WAS test. The clinical findings of AA were evaluated according to the previously described criteria for human AA (Olsen EA, et al, 2004). KO WAS mice: *cchcr1* ^*-/-*^ mice subjected to the WAS test WT WAS mice: wild-type mice subjected to the WAS test

**Fig 2.**
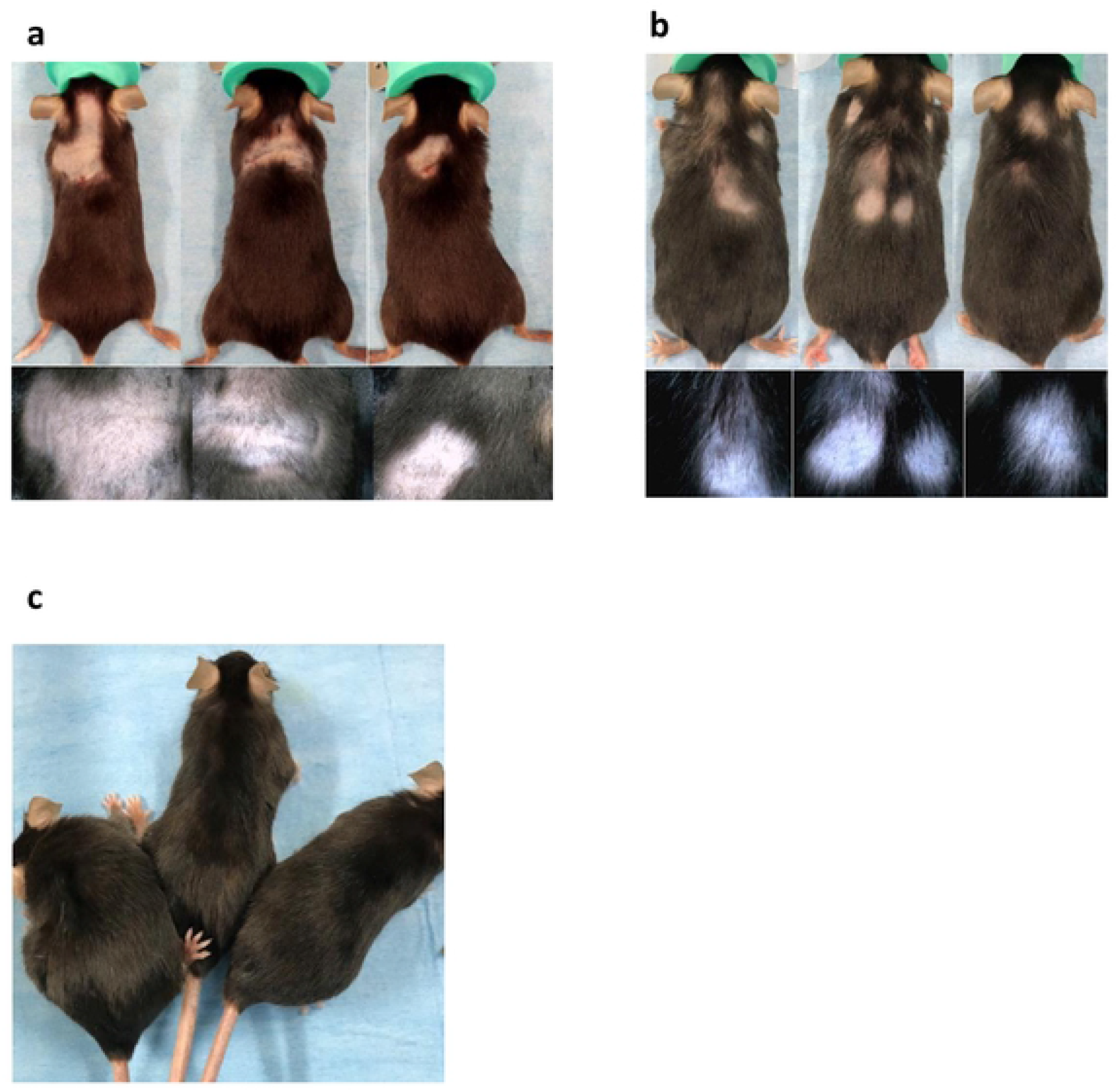
**Fig 2a.** Eight weeks after the WAS test, 25% of *cchcr1*^*-/-*^ mice showed localized hair loss. Short stubs of hair and exclamation mark hairs were seen under dermoscopy. **Fig 2b.** Eighteen weeks after the WAS test on *cchcr1* ^*-/-*^ mice. Recovery from localized hair loss was evident, but the prognosis was variable, and AA-like lesions recurred in some mice. Thin hairs and white hairs were observed under dermoscopy. **Fig 2c.** WT mice after the WAS test.

Eighteen weeks after the WAS test, the localized hair loss spontaneously recovered (Fig. 2b). Regrowth began one to three months after hair loss and was followed by the recurrence of hair loss in the same or other areas. In addition, AA showed a variable prognosis in each mouse and could recur. In aging AA mice, intralesional hair regrowth was a highly characteristic feature of AA; the newly sprouted hairs were white and became thinner as they gradually approached the dorsum, and the majority of the hair follicles were catagen and telogen follicles.

The AA-like skin lesions in *cchcr1* ^*-/-*^ mice induced by WAS were tested by HE staining, immunohistochemistry and scanning and transmission electron microscopy (SEM and TEM). The histopathologic appearance of AA mice included a variety of features. The follicles showed little inflammatory lymphocytic infiltration in the peribulbar region (Fig. 3a). We also observed the infiltration of plasma cells and an increase in the number of hair follicles as well as miniaturized anagen follicles. The bulbs of the hair follicles were infiltrated by CD4- and CD8-positive T cells. On the other hand, there was no infiltration of CD4- and CD8-positive T cells in the specimens from wild-type mice (Fig. 3b).

**Fig 3.**
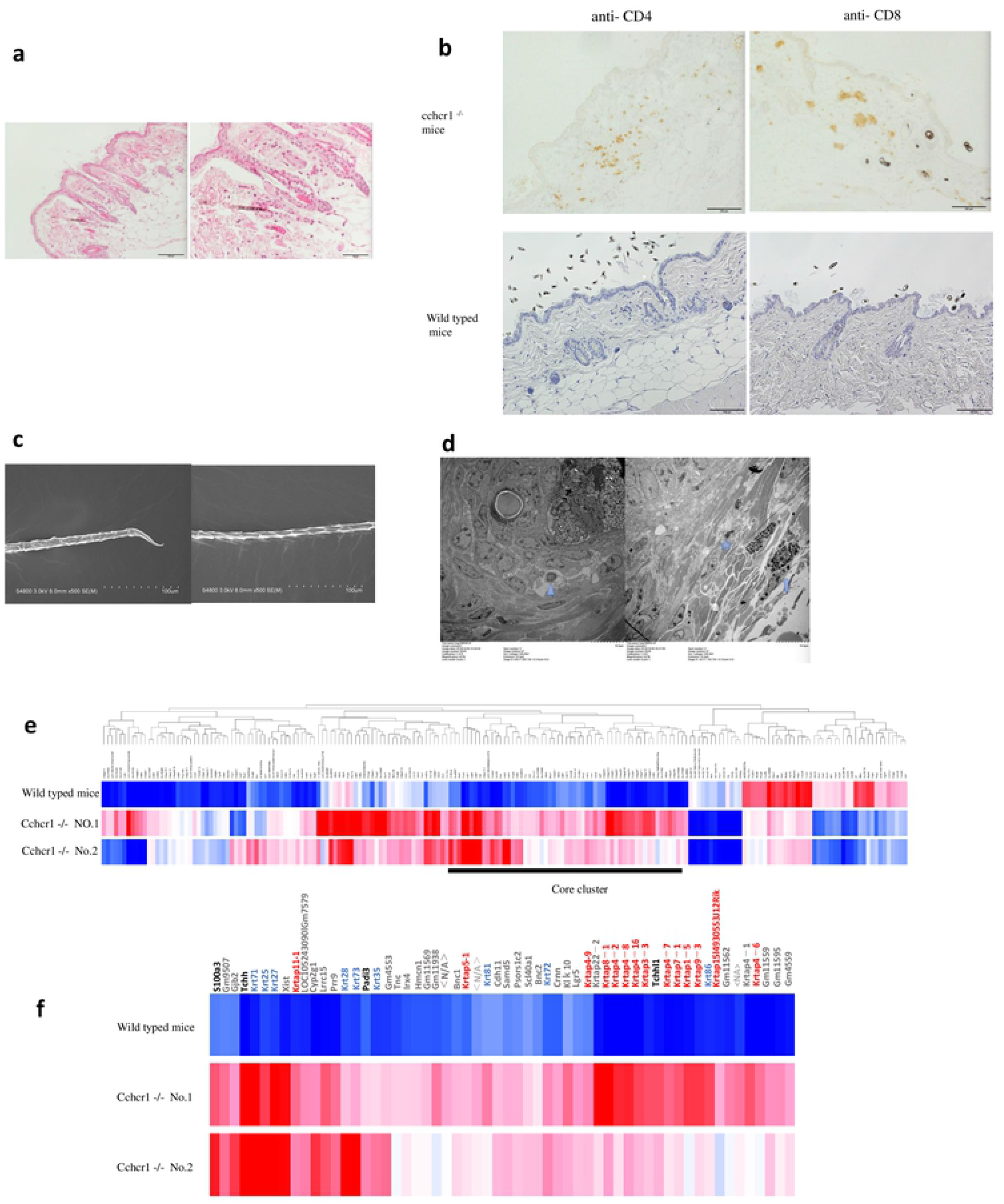
**Fig 3a.** Histological analysis of alopecic lesions in *cchcr1* ^*-/-*^ mice using HE staining revealed follicles with little inflammatory lymphocytic infiltration in the peribulbar region (HE staining; scale bar=100 µm, 50 µm). **Fig 3b.** IHC staining of the skin using CD4+ and CD8+ antibodies showed peribulbar lymphocytic inflammation in alopecic lesions in *cchcr1* ^*-/-*^ mice (IHC stain; scale bar=100 µm). **Fig 3c.** SEM examination of the hair shafts of *cchcr1* ^*-/-*^ mice. (a) Exclamation mark hairs. (b) Abnormal hair shaft formation (Scale bar=100 µm). **Fig 3d.** TEM examination of the skin from alopecic lesions on *cchcr1* ^*-/-*^ mice revealed lymphocytes, dark cells, and mast cells in a perifollicular location (left panel). Apoptosis in the outer root sheath of catagen follicles, matrix keratinocytes and anagen hair bulbs was observed (right panel) (Scale bar=10 µm). **Fig 3e.** Expression analysis of *cchcr1* ^*-/-*^ mice. Heat map of 196 probes showing ≥ 4-fold change in gene expression. **Fig 3f.** Heat map of core cluster genes. The color code depicts KRTAP family (red), keratin family (blue), other hair-related (black), and nonhair-related (gray) genes.

SEM examination of skin from *cchcr1* ^*-/-*^ mice revealed exclamation mark hairs and hair shaft abnormalities as well as the loss of the cuticle layer and exposure of the underlying cortex (Fig. 3c). Dark cells and mast cells in a perifollicular location were observed by TEM examination (Fig. 3d).

To investigate the biological functioning underlying the observed hair loss in cchcr1-/- mice, we performed gene expression microarray analysis of dorsal skin biopsies of cchcr1-/- mice and compared them with that of wild-type mice as a control. This analysis identified 196 genes with 4-fold or greater up- or downregulation. Cluster analysis of the probes revealed that the upregulated gene cluster in cchcr1-/- mice included hair-related genes, such as the keratin and KRTAP genes (Fig. 3e, 3f). Hair keratins and KRTAPs are the major structural components of the hair shaft and are specifically expressed in the medulla, cortex, and cuticle. Other upregulated genes included S100 calcium binding protein A3 (S100A3, substrate of PADI3), trichohyalin (Tchh, substrate of PADI3), peptidyl arginine deaminase type III (Padi 3, related to posttranslational modification) and homeobox C13 (Hoxc13, related to hair shaft differentiation). This result suggested that cchcr1 affects the hair shaft and its formation at the genetic level.

## Discussion

We previously reported genome-edited mice with a nonsynonymous variant of the *CCHCR1* gene in the AA risk haplotype [6]. Among these mice, 2 (12.5%) of the 16 heterozygous mice and 15 (55.5%) of 27 homozygous mice showed patchy alopecia after birth without any stimulation (assessed for up to 10 months). Both the genome-edited mice and AA patients with the risk allele displayed morphologically impaired hair growth and comparable differential expression of hair-related genes, including genes encoding hair keratin and keratin-associated proteins (KRTAPs) [6]. These results were consistent with the microarray analysis in this study; the upregulation of keratins, the KRTAP gene, and the Padi3, S100A3 and Hoxc13 genes was confirmed (Fig. 3e, 3f), suggesting that cchcr1 deficiency caused very similar gene expression patterns in humans and mice with AA carrying risk variants in CCHCR1. In addition, in this study, SEM examination of hair from *cchcr1* ^*-/-*^ mice revealed exclamation mark hairs and hair shaft abnormalities. These findings strongly supported the hypothesis that deficiency or dysfunction of *CCHCR1* is related not only to the hair loss phenotype but also to hair keratinization abnormalities.

Another finding that should be discussed is the presence of peribulbar lymphocytic infiltration in biopsy samples from human AA and mouse AA models because this phenomenon is thought to be one of the markers of AA diagnosis. Peribulbar lymphocytic infiltration was obvious in the alopecic area of *cchcr1* ^-/-^ mice (Fig. 3a, 3b), but it was not observed in genome-edited mice. It is possible that the WAS test could induce peribulbar lymphocytic infiltration in *cchcr1* ^-/-^ mice, but another possibility is that lymphocytic infiltration is not always observable in all alopecic skin lesions. In fact, Müller et al. reported that peribulbar lymphocytic infiltration and lymphocytic infiltration within the fibrous streamers could be seen in only 39% of horizontal sections of human AA skin samples [10].

*CCHCR1* is located 110 kb away from the HLA-C locus and is positioned between the *CDSN* and *SC1* genes [11], and it has also been thought to be involved in the pathogenesis of psoriasis in a transgenic mouse model [12]. Our study supported the involvement of *CCHCR1* in AA pathogenesis, but further experiments will be needed to prove the involvement of *CCHCR1* in psoriasis.

Additionally, past studies indicated that psychological stress might be a triggering factor in AA patients [7]. Therefore, to induce AA-like lesions in *cchcr1* ^-/-^ mice, we utilized the “mild form” of the WAS test in this study. Briefly, the WAS test was performed for only two hours a day and was repeated five times per week for two weeks. The stress sessions were performed between 1000 and 1300 hours to minimize the effect of the circadian rhythm [13]. Nonetheless, 25% of *cchcr1* ^-/-^ mice spontaneously exhibited noninflammatory foci of alopecia on the dorsal side. Grooming and barbering, which are well-known behaviors among stressed C57BL/6 mice, were not observed. The hair loss site was located on the back, not the face, which resulted in characteristic barbering behavior. The morbidity rate of *cchcr1* ^-/-^ mice is higher than that (20%) of C3H/HeJ mice, which is a well-known model for AA [14]. This result strongly indicates that not only the WAS test, which is a psychological stress test, but also genetic factors (i.e., deficiency or nonsynonymous variants of *CCHCR1*) might be responsible for AA. This model also revealed unique clinical findings regarding the female predisposition toward AA (Table 1). A previous report indicated that the relative incidence of AA in one production colony of C3H/HeJ mice was 0.25% for female mice and 0.035% for male mice, but selective breeding raised the frequency to nearly 20% [14]. Aging C3H/Hej mice are known to exhibit alopecia spontaneously. However, in this study, *cchcr1*^-/-^ mice that underwent the WAS test developed hair loss at approximately 16 weeks of age. Therefore, the WAS test could trigger AA in nonelderly mice. The unequal induction of hair loss in female and male mice suggested that the AA that developed in mice shares many features with AA in humans.

In conclusion, we demonstrated for the first time that in *cchcr1* gene knockout mice on the C57BL/6N background, psychological stress can trigger AA-like lesions. Mice deficient in *cchcr1* are likely to be a good model to examine the pathophysiology of AA. The new AA mouse model enables us not only to further investigate AA pathogenesis but also to explore new therapeutic strategies and test the therapeutic efficacy of a wide range of candidates in preclinical studies.

## Materials and methods

### Generation of cchcr^-/-^ mice

The targeting vector was constructed by insertion of loxP sequences within exon 4 of the cchcr1 gene. Exon 4 was fused to cDNA fragments encoded by exons 2 and 3 and exons 5-10. We electroporated the targeting vector into mouse RENKA ES cells (C57BL/6N), selected cells with Geneticin (Invitrogen, Carlsbad, CA), and then screened the cells for homologous recombinants with PCR using the following primer set: sc_*5AF2*: 5’-CAA CTG AGG GCC GTT ACA GAG-3’ and *neo_108r*: 5’-CTT CAG AAG AAC TCG TCA AGA AG-3’. Homologous recombination of these clones was also confirmed by genomic Southern hybridization by probing with a neomycin resistance gene. Genotyping of mice via PCR was performed using the following two primers: *5A-F2*: 5’-CTC ACC TAG AAT TCA GAC ATC C-3’ and *3A-R1*: 5’-AGT TGC AAC TGG CTA TAG CTG C-3’ (shown in Fig. 1b). *Cchcr1* knockout mouse production was performed by TranGenic Inc. (Fukuoka, Japan) in accordance with institutional guidelines. For the control, age-matched C57BL/6N mice were used. Mice were housed in specific pathogen-free facilities, and all the mice were maintained in an environmentally controlled room (illuminated between 08:00 and 20:00) and were fed a pelleted laboratory diet, with tap water available ad libitum. The experimental protocol was approved by the Ethics Review Committee for Animal Experimentation of Juntendo University.

### RT-PCR analysis

cDNA was synthesized from 5 μg of DNase I–treated total RNA using Applied Biosystems 7500 Real-Time PCR Software (Life Technologies Co., Carlsbad, CA, USA). The specific primers for each gene were as follows: 5’-GCA GAA GAT GAG GTT GGA GAC T -3’ and 5’-CTT GTG CTT CGC CTC TTC CA -3’ for *CCHCR1* and 5’-TCC TTC TTG GGT ATG GAA TCC TG -3’ and 5’-GAG GTC TTT ACG GAT GTC AAC G-3’ for β-actin.

### Western blotting analysis

Protein from skin tissue was suspended in a buffer containing 150 mM NaCl, 50 mM Tris-HCl (pH 7.2), 10% ethylene glycol tetraacetic acid, and 1% protease inhibitors (Sigma). The suspension was extensively homogenized with a motor-driven glass/Teflon homogenizer. The resultant homogenate was mixed with SDS-PAGE sample buffer and incubated at 100 °C for 5 min. Western blotting analysis was performed as described previously. Actin was used as the loading control.

#### 2.4. WAS test

The stress procedure was performed according to previous methods, with slight modifications [9]. Mice were placed on a platform (10×10×8 cm) attached to the bottom of a plastic tank (45 cm length × 25 cm width × 25 cm height). The tank was filled with warm water (25 °C) to within 1 cm of the top of the block. The mice were placed on the platform to avoid the water stimulus for two hours, and the test was repeated on weekdays for two weeks.

### Morphological analysis

Macroscopic findings and dermoscopic images were observed regularly. To prepare the sections for histological analysis, AA skin samples were fixed in formalin solution and embedded in paraffin. The sections were stained with hematoxylin and eosin (H&E). For immunohistochemical analysis, the following primary antibodies were used: anti-rabbit CD4 (ab183685, Abcam, Cambridge, MA, USA) and biotin anti-mouse CD8a (BioLegend, San Diego, CA, USA). Tissue sections were counterstained with hematoxylin.

### Electron microscopy

For transmission electron microscopy, the skin samples were fixed in Karnovsky’s fixative and embedded in epoxy resin according to standard procedures. Ultrathin sections were prepared, stained with uranyl acetate and lead citrate, and observed with an HT7700 (HITACHI, Tokyo, Japan) transmission electron microscope. For scanning electron microscopy observation, the hair shafts were dehydrated in 100 % ethanol. After coating with platinum, the samples were examined with an S-4800 field-emission scanning electron microscope (Hitachi, Tokyo, Japan).

### Total RNA Extraction

Frozen mouse skin tissue was homogenized with TRIzol reagent (Thermo Fisher Scientific, Inc., Waltham, MA, USA) using zirconia beads. The tissue lysate was processed according to the manufacturer’s instructions. RNA samples were purified by an RNeasy MinElute kit (QIAGEN N N.V., Venlo, Netherlands).

### Microarray analysis of mouse skin RNA

One hundred nanograms of total RNA was processed for use on the microarray using the GeneChip WT PLUS Reagent Kit (Thermo Fisher Scientific, Inc., Waltham, MA, USA) according to the manufacturer’s instructions. The resultant single-stranded cDNA was fragmented and labeled with biotin and then hybridized to the GeneChip Mouse Gene 2.0 ST Array. The arrays were washed, stained and scanned using the Affymetrix 450 Fluidics Station and GeneChip Scanner 3000 7G (Thermo Fisher Scientific, Inc.) according to the manufacturer’s recommendations. The expression values were generated using Expression Console software, version 1.4 (Thermo Fisher Scientific, Inc.) with the default robust multichip analysis parameters.

## Acknowledgments

This work was supported in part by a Grant-in-Aid for Scientific Research from the Ministry of Education, Culture, Sports, Science and Technology of Japan (AO or SI). We thank the members of the Laboratory of Morphology and Image Analysis, Research Support Center, Juntendo University Graduate School of Medicine, for technical assistance.

## Author Contributions

### Investigation

Qiao-Feng Zhao, Nagisa Yoshihara, Atsushi Takagi, Shigaku Ikeda Writing original draft: Qiao-Feng Zhao, Nagisa Yoshihara, Shigaku Ikeda Writing review & editing: Nagisa Yoshihara, Shigaku Ikeda

### Conceptualization

Nagisa Yoshihara, Shigaku Ikeda Data curation: Qiao-Feng Zhao, Atsushi Takagi Formal analysis: Nagisa Yoshihara, Shigaku Ikeda Funding acquisition: Shigaku Ikeda

### Methodology

Qiao-Feng Zhao, Nagisa Yoshihara, Atsushi Takagi, Etsuko Komiyama, Akira Oka, Shigaku Ikeda

### Resources

Shigaku Ikeda Validation: Shigaku Ikeda

